# Endocannabinoid-mediated rescue of somatosensory cortex activity, plasticity and related behaviors following an early in life concussion

**DOI:** 10.1101/2024.01.30.577914

**Authors:** J. Badaut, L. Hippauf, M. Malinconi, B.P. Noarbe, A. Obenaus, C. J. Dubois

## Abstract

Due to the assumed plasticity of immature brain, early in life brain alterations are thought to lead to better recoveries in comparison to the mature brain. Despite clinical needs, how neuronal networks and associated behaviors are affected by early in life brain stresses, such as pediatric concussions, have been overlooked. Here we provide first evidence in mice that a single early in life concussion durably increases neuronal activity in the somatosensory cortex into adulthood, disrupting neuronal integration while the animal is performing sensory-related tasks. This represents a previously unappreciated clinically relevant mechanism for the impairment of sensory-related behavior performance. Furthermore, we demonstrate that pharmacological modulation of the endocannabinoid system a year post-concussion is well-suited to rescue neuronal activity and plasticity, and to normalize sensory-related behavioral performance, addressing the fundamental question of whether a treatment is still possible once post-concussive symptoms have developed, a time-window compatible with clinical treatment.

## Introduction

A conserved evolutionary aspect of all living creatures is the necessity to adapt its behaviors according to changes in the environment. In complex animals, treatment of the sensory inputs that relate to the environment require proper integration and processing of the stimuli to ensure adaptative response and survival. In a number of brain disorders, the central processing of sensory inputs is altered, leading eventually to the overall morbidity of the condition. In human subjects, failure to properly integrate sensory inputs leads to a wide variety of impairments, considerably worsening the patient’s quality of life. These alterations can occur at any time during the lifespan, and an often-overlooked question, yet essential to develop *ad hoc* therapeutical strategies, is whether these alterations would lead to the same consequences in the developing brain in comparison to the adult brain. Indeed, because of the on-going plasticity mechanisms in the early times of brain development, it is often assumed that this would compensate for the disturbances^1^.

While brain injuries can lead to focal disruption of specific brain regions, diffuse injuries, typically seen in concussions, initiate a brain-wide neurometabolic cascade that have the potential to durably disrupt brain functioning. While for concussions recovery will be achieved within days, post-concussion symptoms might perdure months to years, for 10 to 20% of adults^2,3^. While all age groups are likely to be affected by head concussion, children and adolescents represent a large fraction of emergency admissions for concussions to mild traumatic brain injuries (mTBI), and are more susceptible to developing persistent post-concussive syndromes^4,5^. Those symptoms include cognitive, psychosocial and physical deficits with a high prevalence in altered sensory integration such that the assessment of balance and postural stability (that are likely the result of central sensory integration dysfunction rather than vestibular or oculomotor alterations^6^) are a means to assess the extent of the impairment and its recovery. Considering this higher susceptibility to early life events, it is confounding that the underlying cellular and molecular mechanisms have been overlooked in juvenile models of concussion, especially in the long-term range. Without that knowledge, specific treatments are currently unavailable in clinical settings.

Sensory processing involves various brain regions, with the somatosensory cortex representing a critical hub receiving sensory neuronal inputs. In several adult models of moderate to severe TBI, the neuronal activity of the somatosensory cortex has been shown to be exacerbated acutely following the ipsilateral injury^7–9^. In those models, neuronal activity of other brain regions has also been shown to be altered^8–15^. Similar consequences were found within weeks following a juvenile moderate to severe TBIs in the somatosensory cortex^16^ as well as in the hippocampus^17^. While understanding the full time-course of concussion -induced pathology is critical to delineate efficient therapies and despite evidence of the long-term alterations even after mTBI^18^, the gap in knowledge of the mid to long term (*i.e.* 3 months or more after injury) functional consequences of TBIs remains to be elucidated with a chronic functional evaluation from neuronal activity to behavioral outcomes .

As noted there are still no treatments for the consequences of TBI. Endocannabinoids are lipid mediators that exert a fine tuning of synaptic homeostasis^19–22^ and exhibit anti-inflammatory properties ^23,24^. A critical feature of this neuromodulatory system is an on-demand synthesis and release^25^ (*i.e.* as a function of cellular activity). As endocannabinoids are also tightly regulated by their degradation, pharmacological inhibition of endocannabinoid degradative enzymes has been proposed as a potential therapeutic approach for the treatment of TBIs ^10–12,26,27^. Nonetheless, therapeutic preclinical studies mainly during the early phase of primary injury (*i.e.* minutes after injury), relied on the anti-inflammatory properties of endocannabinoids. In clinical practice, patients are rarely seen in that time-window, especially considering concussions/mTBIs, when patients seek care only after the development of post-concussive symptoms. This left unanswered the critical question of whether treatment is still possible in the later chronic phases of injury.

Here we show in a pediatric mouse model that a single concussion triggers long lasting consequences to sensory integration in somatosensory cortex and alters associated behaviors. This is due to a higher level of neuronal activity in GABAergic neurons of the somatosensory cortex that blunts behaviorally-induced sensory integration. We also provide evidence that pharmacological intervention is still possible when symptoms are already set in the time and that the endocannabinoid system could represent an innovative approach for treatment of long-term post-concussive symptoms.

## Results

### Consequences of an early life concussion on somatosensory cortex activity, plasticity and related behaviors

Long-term consequences of an early in life brain alteration on neuronal activity were assessed using the Closed-Head Injury with Long-term Disorders (CHILD) model^28^ that exhibits behavioral and morphological alterations up to 12 months^18,29^. Mice received a single mild impact at P17 in the CHILD group (CHILD group, n=4) while sham mice underwent the same procedure without impact (n=5, Fig.1). We next induced a month later the expression of the calcium indicator GCaMP6f in GABAergic neurons from layers 2/3 and 4 of the somatosensory cortex (SSC, see methods, Fig. 1, Fig 2d) and implanted through a gradient refractive index (GRIN) prism (Fig.1). The correct location in the SSC of the implants was later confirmed from post-mortem in MRI scans (Extended data Fig. 1).

**Figure 1.**
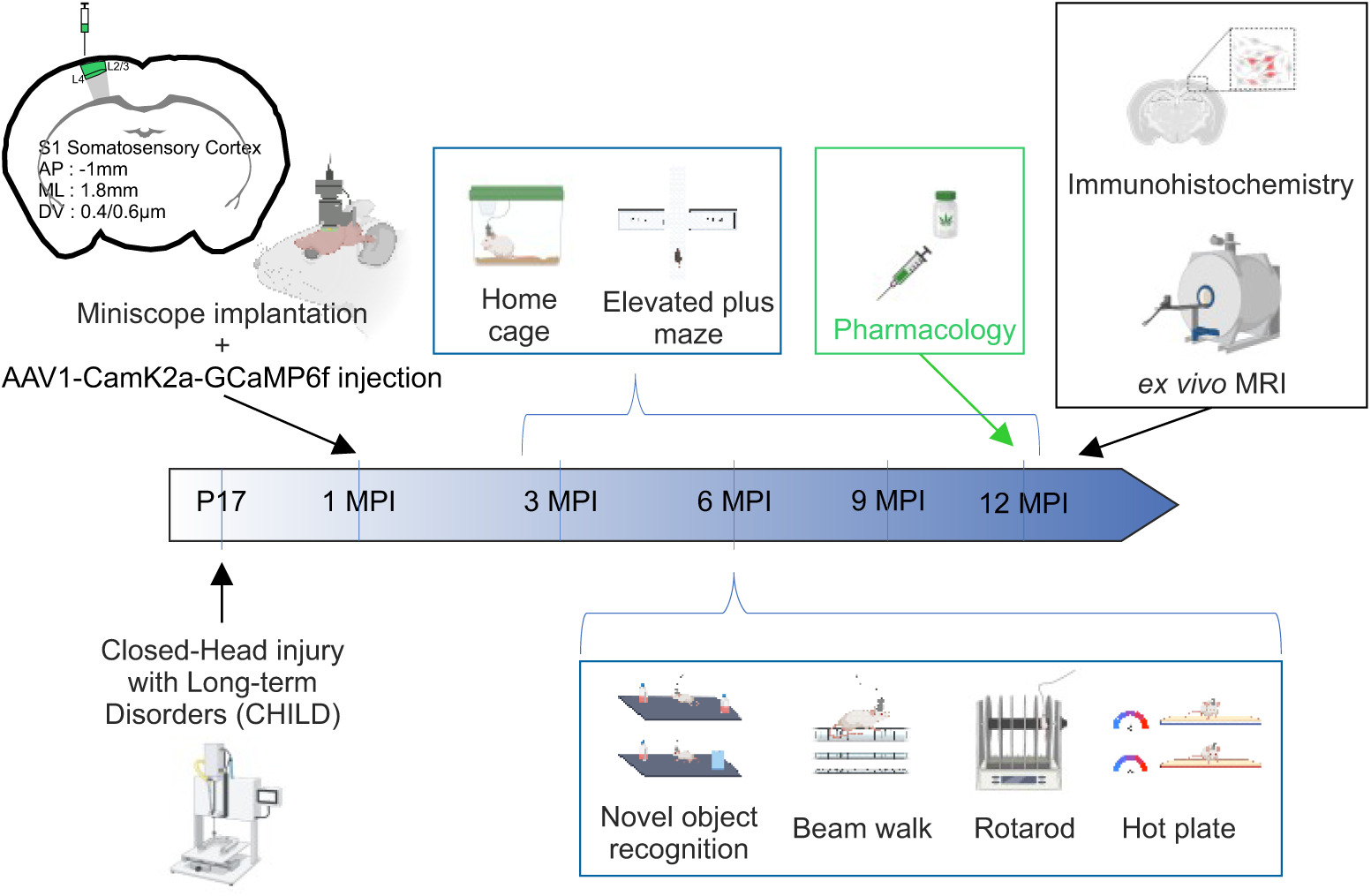
Experimental protocol. Seventeen days after birth, mice from the CHILD group received a single mild traumatic brain injury, while animals from the sham group followed the same procedure at the exception of the impact. A month later, injections of a virus allowing the expression of GCamp6F under the CamK2a promoter were performed in the somatosensory cortex of all mice and the GRIN lens were implanted at the same location. Every 3 months following TBI, neuronal somatosensory cortex activity of all animals was evaluated without significant sensory stimulation (in a typical home-cage) or while on an elevated plus maze. At the 6-month post-injury time point, animals were also probed with the novel object recognition task, beam walk task, rotarod and hot plate test. All these tests were designed to focus on sensory integration. One year after concussion, the effects on endocannabinoid degradation inhibition were pharmacologically tested in a home-cage and elevated plus maze. *Ex vivo* MRI and immunohistochemistry experiments were then performed.

**Figure 2.**
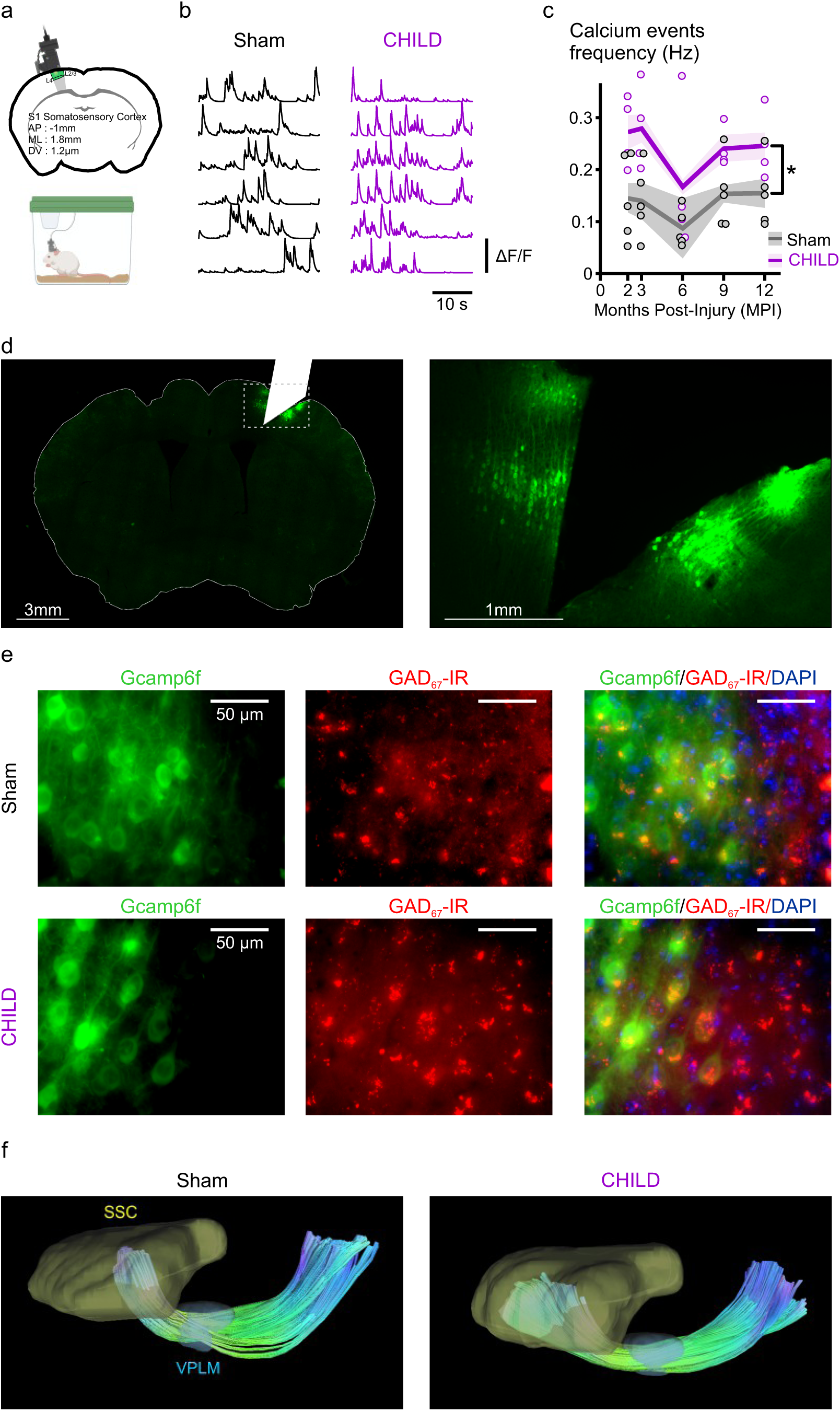
Consequences of an early life concussion on somatosensory cortex activity and imaging. **a.** While in a typical home-cage, calcium imaging in the somatosensory cortex was performed *in vivo* in unanesthetized mice. **b.** Representative ΔF/F calcium recordings from sham and CHILD mice. **c.** Twelve-month time course of calcium events frequency in somatosensory cortical neurons (Hz) in sham (n=5) and CHILD mice (n=4). CHILD mice exhibited a significant increase in activity in those neurons regardless of the time point (2-way RM ANOVA, *F*_(1,28)_=8.064, *p*=0.025). **d.** Typical Gcamp6f fluorescence at low (left) and high (right) magnifications. The white notch represents the GRIN lens location. **e.** representative fluorescence images of Gcamp6f and GAD_67_-immunofluorescence in sham (top) and CHILD (bottom) animals. GAD_67_-immunofluorescence was found in 99% of the sham neurons evaluated (158/159, in 3 animals) and in 94% of the CHILD neurons (147/156, in 3 animals). **f.** DTI tractography mapped from the left medial ventral posterior lateral nucleus (VPLM) of the thalamus and terminating in the somatosensory cortex (SSC). As can be visualized, CHILD mice tended to exhibit lower density fiber (n=3 for both groups, p>0.05, see extended data table 1), an increased tract dispersion between the VPLM and the SSC (n=3 for both groups, p>0.05, see extended data table 1) and a shallower entry angle in the SSC (n=3 for both groups, p>0.05, see extended data table 1). Average values are shown as solid lines, SEM is represented as the shaded area. *, *p*<0.05.

With this model we first investigated the consequences of a single concussion on neuronal activity in an empty housing-type cage, in absence of significant sensory stimulation (Fig. 2a). Two months post injury (MPI), CHILD mice exhibited a significant 88 ± 20% increase in calcium-related events frequency (unpaired t-test, *p*<0.05) when compared to their sham controls (Fig. 2b, c). Strikingly, this increased activity in the somatosensory cortical neurons persisted up to a year post injury (2-way RM ANOVA, *F* _(1,28)_= 8,064, *p*<0.05). Post-mortem immunohistochemistry analysis revealed that 99% and 94% of the neurons expressing GCamP6f in sham and CHILD mice respectively, colocalized with GAD67 immunolabelling (Fig. 2e), suggesting that the vast majority of transfected neurons were GABAergic. The cortico-thalamo-cortical network plays a critical role in the flow of information between the thalamus and sensory areas of the cortex^30^. Thalamocortical projections undergo structural reorganization without laminar-specific targeting following TBI in more mature or severe preclinical models^31–33^. Tractography analysis was therefore performed from *ex vivo* diffusion-MRI for each group at 12 months after concussion. Overall, and in accordance with previous reports from TBI models^31^, thalamocortical projections tended to be less dense (-17%) and more widespread within the SSC after a single concussion (sham span of 21 ± 1 mm, CHILD span of 24 ± 1 mm, n=3 per group, unpaired t-test, *p*=0.098, Extended data table 1). In addition, analysis found that thalamocortical fibers were predominantly oriented with an angle with the pial surface of the SSC (65 ± 6°, n=3), this was shallower in CHILD brains (54 ± 3°, n=3, Fig. 2f). These altered cortical projections have been previously linked to hyperactivity of the thalamo-cortical networks^31^, suggesting a potential mechanism.

Mice were then challenged in the elevated plus maze at 3 months after injury (Fig. 3a). While sham mice spent more time in closed arms (163 ± 27 seconds) rather than open ones (78 ± 20 seconds, paired t-test, *p*=0.031). After a single concussion, mice spent an equal amount of time in both types of arms (118 ± 17 seconds in closed arms and 114 ± 17 seconds in open arms, paired t-test, *p*=0.995, Fig. 3b). While a differential preference for the type of arms in the elevated plus maze is usually interpreted as a change in the anxiety state of the animal, a lack of discrimination could as well represent the hallmark of altered sensory integration after an early in life concussion. When focusing on the z-score of calcium-related fluorescence of individual cells, the entry into the open arms in sham mice was associated with increased cell fluorescence across the entire SSC depth (Fig. 3c). Accordingly, overall SSC neuronal activity exhibited a transient and significant increase in the frequency of calcium-related events at the time of the entrance of the sham animal in open arms (Fig. 3d). This transient rise in SSC activity appeared to be important for the detection of the change in environment, as its amplitude was negatively correlated to the number of entries in the open arms made by the animals (linear regression by Pearson’s correlation, *p*<0.05, Fig. 3e, see also Fig 4i and Extended data Fig 3c). In sharp contrast, we found no change of fluorescence (Fig. 3c) or cortical neuronal activity (Fig. 3d) in mice from the CHILD group, suggesting the absence of sensory integration upon entrance into open arms. This lack of cortical plasticity induced by the change in environment in CHILD animals could therefore account for the higher of number of entries in those arms compared to the sham group. Altogether our results indicate that a single early in life concussion triggers a long term (*i.e.* at least up to a year post-injury) cortical hyperactivity (Fig. 2c) and altered behaviorally-induced cortical plasticity (Fig. 3f, 2-way RM ANOVA, sham *vs* CHILD regardless of the time point, *F* _(1,23)_ =21.309, *p*=0.004). Those alterations in the SSC are also directly associated to a change in the behavior of the animals.

**Figure 3.**
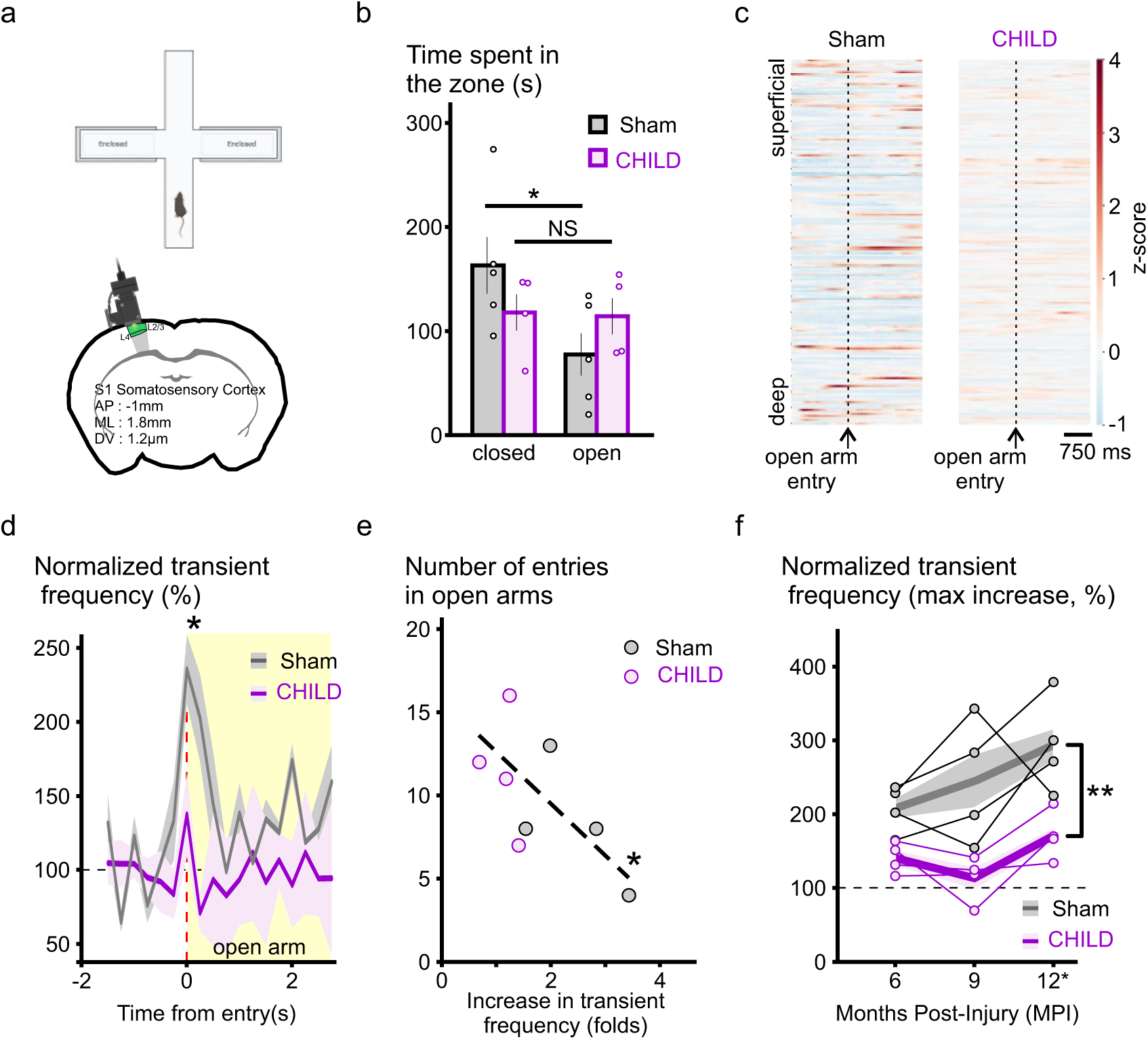
Consequences of an early life concussion on somatosensory cortex plasticity induced by sensory integration in the elevated plus maze. **a.** While on an elevated plus maze, calcium imaging in the somatosensory cortex was performed *in vivo* in unanesthetized mice. **b.** Group data showing the time spent in open and closed arms of the elevated plus maze for sham (n=5) et CHILD (n=4) mice. While sham mice spent more time in the closed arms (163 ± 27 s vs 78 ± 20 s in open arms, paired t-test, *p*=0.031), CHILD spent indistinctly the same amount of time in each type of arm (open arms 114 ± 17s, closed arms 118 ± 17 s, paired t-test, p=0.995). **c.** z-score plots of representative sham and CHILD animals with cells ordered in function of their depth in the somatosensory cortex (with the deepest cells at the bottom). Plots are averages of 3 to 5 events and centered to the entrance of the animals into the open arms of the maze (dashed lines). Increased neuronal activation at the entrance is seen in sham animals (in red, after the dash line) but not in CHILD mice. **d.** Average plots of the averaged normalized calcium-transients frequency of sham (grey, n=5) and CHILD (purple, n=4) groups. A significant increase in neuronal calcium activity occurred at the entrance in the open arms (red dash line) in sham mice (+136 ± 23%, paired t-test, *p*= 0.039) but was blunted in CHILD mice (+37 ± 28%, paired t-test, *p*=0.747). **e.** Scatter plot of the number entries in the open arms plotted *vs* the amplitude of the increase in calcium-transients frequency in sham (grey, n=5) and CHILD (purple, n=4) mice. Overall an inverse relationship was found between these 2 parameters (Pearson’s correlation, *p*=0.049, see also panel i in figure 4). **f.** Twelve-month time course of calcium events frequency increase at the entrance in the open arms in somatosensory cortical neurons in sham (n=5) and CHILD mice (n=4). CHILD mice exhibited a blunting of this form of neuronal sensory integration in those neurons regardless of the time point (2-way RM ANOVA, *F*_(1,23)_=21.309, *p*=0.004). Average values are shown as solid lines, SEM is represented as the shaded area (error bars in b.). NS, *p*>0.05, *, *p*<0.05, ** *p*<0.01.

**Figure 4.**
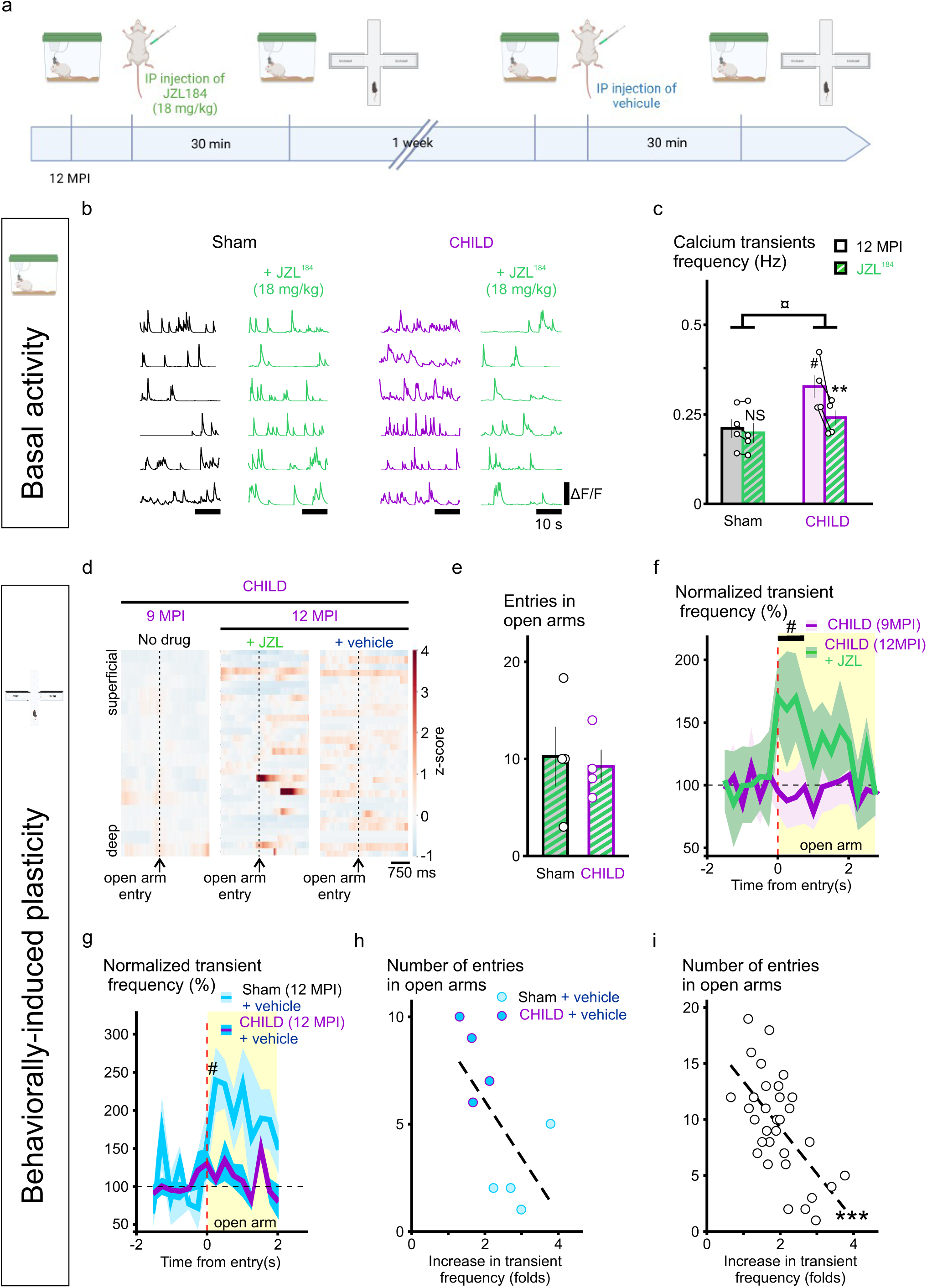
Endocannabinoid-mediated mitigation of the long-term effects of an early life concussion. **a.** Experimental procedure for the pharmacological intervention 12 months after the TBI using a specific inhibitor of endocannabinoid degradation induced by the monoacylglycerol lipase (JZL^184^, 18 mg/kg). **b.** and **c.**, calcium-related neuronal activity in the somatosensory cortex recorded in a typical home-cage, 30 minutes after an IP injection of JZL^184^. **b.** Representative ΔF/F calcium recordings from sham and CHILD mice, before (left, black and purple traces, respectively) and 30 minutes following the IP injection of JZL^184^ (right, green traces). **c.** Group data showing that the potentiation of endocannabinoid signaling by JZL^184^ injection specifically targeted basal calcium-related neuronal activity in CHILD mice (2-way RM ANOVA, *F*_(1,15)_=10.444, *p*=0.018, ¤). JZL184 had no effect in sham mice (n=4, tukey posthoc, *p*=0.45, NS) while it reduced calcium related neuronal activity in the somatosensory cortex of CHILD mice (n=4, tukey posthoc, *p*=0.002, **) to sham levels (n=4 in both groups, tukey posthoc, *p*=0.364). **d. to i.**, endocannabinoid-dependent modulation of calcium-related neuronal plasticity induced by the entrance in open arms of the elevated plus maze and associated behavior. **d.** z-score plots of a representative CHILD mouse 9 MPI, and 12 MPI 30 min after JZL184 injection and 12MPI 30 min after vehicle injection, with cells ordered in function of their depth in the somatosensory cortex (with the deepest cells at the bottom). Plots are averages of 3 to 5 events and centered to the entrance of the animals into the open arms of the maze (dashed lines). Increased neuronal activation at the entrance is seen when the animal was injected with JZL^184^ (in red, center plot after the dash line) but not at 9MPI or when injected with a vehicle solution. **e.** Group data of the number of entries in the open arm by sham (n=4) and CHILD (n=4) mice 30 min after injection. Both groups exhibited the same behavior (unpaired t-test, *p*=0.814). **f.** Average plots of the averaged normalized calcium-transients frequency of CHILD mice at 9MPI (purple, n=4) and at 12 MPI 30 min following a JZL184 injection (green, n=4). A significant increase in neuronal calcium-related activity occurred in CHILD mice (n=4) at the entrance in the open arms (red dash line) in presence of JZL^184^ (+99 ± 9%, paired t-test, *p*= 0.007, #) but not in presence of vehicle (+13 ± 9%, paired t-test, *p*=0.499). **g.** Average plots of the averaged normalized calcium-transients frequency of sham (n=4) and CHILD (n=4) mice at 12 MPI following a vehicle injection. Similarly to the 6 and 9 MPI time points without injection, a significant increase in neuronal calcium-related activity was seen in sham mice (n=4) at the entrance in the open arms (red dash line) after vehicle injection (+106 ± 28%, paired t-test, *p*= 0.012, #) but not in CHILD mice (n=4, +20 ± 13%, paired t-test, *p*=0.194). **h.** and **i.**, Scatter plots of the number entries in the open arms plotted *vs* the amplitude of the increase in calcium-transients frequency at the time of entry. **h.** in sham and CHILD mice 30 minutes after a vehicle injection (sham, light blue n=4; CHILD, dark blue n=4), and **i.** across all the experiments performed from both groups (n=34). Overall a significant inverse relationship was found between the amplitude of calcium-related neuronal activity and the number of entries in the open arms (Pearson’s correlation, *p*<0.01, ***). Average values are shown as solid lines in **f.** and **g.**, SEM is represented as the shaded area (error bars in **c.** and **e.**).

### Endocannabinoid-mediated mitigation of the long-term effects of an early concussion

Such an increase of the neuronal activity of the somatosensory cortex after an early concussion is predicted to prevent further increases in the activity, consequently blunting sensory integration at the time of sensory stimulation. To test this hypothesis, we next investigated the possibility of reducing the spontaneous neuronal activity to rescue behaviorally-induced plasticity. The endocannabinoid system is a major neuromodulatory system with anti-inflammatory properties and neuroprotective effects both *in vitro* and *in vivo* ^34,35^ and is widely described in the somatosensory cortex^36–39^. Furthermore, endocannabinoids are released on-demand in an activity-dependent fashion^25^ and can set the level of activity of neuronal networks^21^. From these unique properties we predicted that increasing endocannabinoid levels would reduce neuronal activity in the most active networks (i.e. observed in the CHILD group), while inducing marginal effects on less active neuronal networks (i.e. in the sham group). Therefore, we injected CHILD and sham mice at 12 months post-injury with JZL^184^ (18 mg/kg, IP), a specific monoacylglycerol lipase blocker that inhibits 2-AG degradation for several hours *in vivo*^40^ and therefore increases endocannabinoid availability (Fig. 4a).

Thirty minutes after JZL^184^ injection, the calcium-related neuronal activity of the somatosensory cortex remained unchanged in the home cage in sham animals (Tukey *posthoc* test, *p*>0.05, Fig. 4b, c and Extended data Fig. 4a). In contrast, when injected with JZL^184^, CHILD animals exhibited a decreased calcium-related neuronal activity in the SSC in absence of significant sensory stimulation (Tukey *posthoc*, *p*<0.05), that matched the level of activity of sham animals (tukey *posthoc*, *p*>0.05, Fig. 4b, c and Extended data Fig. 4a).

Hence, in absence of substantial sensory stimulation, JZL^184^ reduced the calcium-related neuronal activity of the somatosensory cortex of CHILD animals but not shams (2-way RM ANOVA, p<0.05, Fig. 4c).

We next tested the consequences of calcium-related neuronal activity normalization in CHILD mice on elevated plus maze-induced neuronal plasticity. We found that in presence of JZL^184^ calcium-related neuronal responses induced by the entry into open arms was rescued in CHILD mice (Fig. 4d and 4f). This was also associated with no difference between groups in the number of entries into the open arms (sham+JZL *vs* CHILD+JZL, unpaired t-test, *p*>0.05, Fig. 4e). JZL benefit was both specific and reversible as vehicle injection a week later failed to reproduce JZL effects in the test (Fig.4g). We also found that neuronal responses observed across all sessions in the somatosensory cortex was highly predictive of the behavioral outcomes (*i.e.* the number of entries in the open arms) in the elevated plus maze (Fig. 4h and 4i). Using a classification algorithm to segregate sham from CHILD animals, based on basal network activity (in the home cage), amplitude of the neuronal network’s response to the entry of the open arms of the elevated plus maze and the number of entries of the animals in the open arms as modalities obtained at 6 and 9 MPI, we were able to correctly classify all the animals during the vehicle injection experiment at 12 MPI (loss of 0.1019 and accuracy of 1). Remarkably, when tested on the JZL injection dataset, the algorithm misclassified all CHILD animals as sham animals (loss of 6.0792 and accuracy of 0.5), further supporting the rescue of the phenotype by the JZL injection. These results strongly support that CHILD effects on neuronal activity, plasticity and associated behaviors that can be reversibly dampened a year after injury with the pharmacological intervention of the specific endocannabinoid degradation inhibitor JZL^184^.

### Behaviorally-induced potentiation of neuronal activity alteration is a hallmark of early in life concussion

We also investigated whether other behaviors associated with cortical somatosensory integration are impacted 6 months after early in life concussion. In the novel object recognition task, exposure to the novel object was shown to induce an increase in the activity of the somatosensory cortex, specifically in layers 2/3^41^. In order to determine whether an early in life traumatic experience would blunt other forms of neuronal potentiation, we tested the ability of mice to exhibit novel object-induced neuronal potentiation. For this purpose, mice were first habituated to the empty open field for 5 minutes and then to 2 identical objects for 5 minutes the following 3 days. On the test day, one of the familiar objects was substituted for a new one. Under these conditions, sham mice exhibited an increased interest for the novel object as expressed by an increased preference index (tukey *posthoc*, *p*<0.05, Fig. 5b). As previously described, neuronal activity in the most superficial layers of the SSC (layers 2/3, Fig. 5a) was specifically increased (top plots of Fig. 5c). In contrast, mice that suffered early in life concussion did not exhibit any particular interest for the novel object (tukey *posthoc*, *p*>0.05, Fig. 5b), which was highly different from the sham population (2-way RM ANOVA, *p*<0.05, Fig. 5b). This absence of interest for the novel object was also associated with an absence of specific neuronal response of the SSC induced by the switch in objects (bottom plots of Fig. 5c).

**Figure 5.**
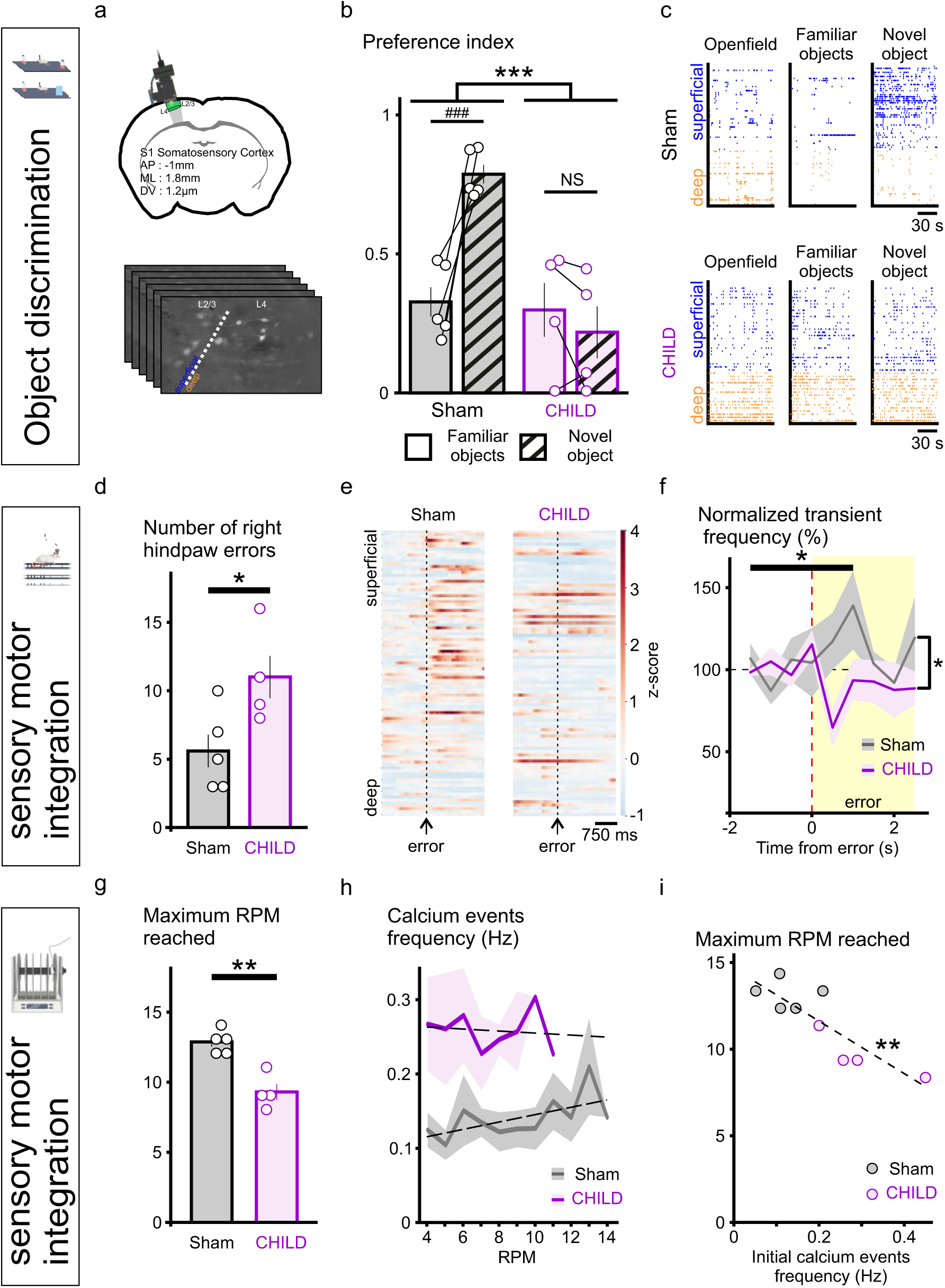
The alteration of behaviorally-induced potentiation of neuronal activity is a hallmark of an early life concussion. **a.**, **b.** and **c. Novel object discrimination task**. **a.** calcium imaging was performed in the left somatosensory cortex (ipsilateral to the injury when applicable). **b.** Group data of the preference index for the novel object at day 5 of the protocol (see methods). Sham (n=5) and CHILD (n=4) mice exhibited differences in attraction to the novel object (2-way RM ANOVA, *F*_(2,23)_=1.155, *p*=0.001, ***). Sham mice spent more time with the novel object (tukey posthoc, *p*<0.001, ###) while CHILD mice did not exhibit preference (tukey posthoc, *p*=0.331, NS). **c.** Whole session representative raster plots of calcium events on day 1 without object (left, ‘openfield’), on day 4 with familiar objects (center, ‘Familiar objects’) and on day 5 when exposed to the novel object recorded in a sham (top) and CHILD (bottom) mice. Cells are organized by their depth in the cortex, with deepest cells at the bottom (orange, layer 4 of the SSC) and most superficial at the top (blue, layers 2/3 of the SSC). When exposed to the novel object sham mice exhibited a specific increase in calcium transients in layers 2/3 of the SSC (top). In contrast CHILD mice exhibited overall higher level of calcium transient frequency in all layers and did not show any change in presence of the novel object (bottom). **d.**, **e.** and **f. Beam walk**. **d.** Group data of the number of errors (paw slips) made by sham (n=5) and CHILD (n=4) mice. CHILD mice made more paw positioning errors compared to sham mice (unpaired t-test, *p*=0.042, *). **e.** z-score plots of representative sham and CHILD animals with cells ordered in function of their depth in the somatosensory cortex (with the deepest cells at the bottom). Plots are averages of 3 to 5 events and centered to a right hindpaw slip (dashed lines). Increased neuronal activation at the error is seen in sham animals (in red, after the dash line) but not in CHILD mice. **f.** Average plots of the averaged normalized calcium-transients frequency of sham (grey, n=5) and CHILD (purple, n=4) mice centered to the mispositioning of the right hindpaw (dashed lines). A significant increase in neuronal calcium-related activity occurred in sham mice at the time of the error (paired t-test, *p*=0.034, *) while this was blunted in CHILD animals (2-way RM ANOVA, *F*_(5,53)_=2.566, *p*=0.044, *). **g.**, **h.** and **i. Rotarod. g.** Group data showing the maximum rod speed sham (n=5) and CHILD (n=4) mice could handle before falling. Sham mice could handle higher speeds (12.8 ± 0.3 RPM) compared to CHILD animals (9.3 ± 0.5 RPM, unpaired t-test, *p*=0.016, *). **h**. Average plot of the calcium-related neuronal activity in GABAergic neurons of the somatosensory cortex according to the speed of the rod. While neuronal activity increased with speed in sham mice (+ 45 % from 0.12 ± 0.02 Hz to 0.18 ± 0.03 Hz at maximum speed, paired t-test, *p*=0.017), it did not significantly increase in CHILD mice (+ 7 % from 0.27 ± 0.06 Hz to 0.28 ± 0.02 Hz at maximum speed, paired t-test, *p*=0.605). i. Plot showing the significant inverse correlation between the calcium-related neuronal activity in the SSC at 4 RPM and the maximum speed of the rod that sham (n=5) and CHILD (n=4) mice could handle (Pearson’s correlation, *p*=0.002, **). Average values are shown as solid lines in **f.** and **h.**, SEM is represented as the shaded area (error bars in **d.** and **g.**).

We next evaluated the consequences of an early in life injury on sensory motor integration in the beam walk task. Overall, sham and CHILD mice performed equally in terms of time to cross 30 cm beams (Extended data Fig. 4c), regardless of the diameter of the beam (2-way RM ANOVA, *p*>0.05, Extended data Fig. 4c), suggesting the absence of major locomotor dysfunctions. However, CHILD mice exhibited an increased number of slips (errors) from of the right hind paws compared to the shams (unpaired t-test, *p*<0.05, Fig. 5d). Remarkably, in sham mice right hind-paw errors were associated with an increase in calcium-related neuronal activity (Fig. 5e and 5f) signifying somatosensory integration. After early in life concussion, mice hind-paw errors did not correlate to a change in calcium-related neuronal activity (Fig. 5e and 5f), suggesting a deficit in sensory processing in the SSC after CHILD (2-way RM ANOVA, *p*<0.05, Fig. 5f). Mice were also tested on the rotarod paradigm to further assess sensory motor integration. Sham mice performed better than CHILD mice, reaching a higher speed on the rotarod before falling (unpaired t-test, *p*<0.05, Fig. 5g). Gradual increase of rotarod speed correlated to increased calcium-related activity in neurons of the SSC in sham group (calcium events frequency at 4RPM *vs* calcium events frequency at max speed in sham mice, paired t-test, *p*<0.05, Fig. 5h). In contrast, the calcium-related activity of the neurons in the SSC did not change with increased rotarod speed in the CHILD group (calcium events frequency at 4RPM *vs* calcium events frequency at max speed in CHILD mice, paired t-test, *p*>0.05, Fig. 5h and Extended data Fig. 5). This further strengthens the hypothesis that mice with early in life concussion were unable to increase neuronal activity in the SSC for proper sensory motor integration (see also Extended data Fig. 5). We also show that regardless of the experimental group the neuronal activity in the SSC at the minimal speed (4 RPM) was highly predictive of the performance of the mice on the rotarod (Pearson’s correlation, *p*<0.05, Fig. 5i).

Our results clearly demonstrate that sensory integration in SSC, and associated behaviors, are dampened after early in life brain concussion when sensory integration is associated to an increase in activity in this cortex.

### Specificity of the alteration of plasticity induced by an early life traumatic brain injury

We next investigated whether a CHILD is also associated with alterations of forms of short-term depression of neuronal activity. We found that sham mice positioned on a slightly heated plate (from 20 to 26°C) for 30 s exhibited a significant overall decrease in calcium-related neuronal activity in the SSC (-73 ± 2% from control activity at 20°C, p<0.001, Fig. 6). Under identical experimental paradigm, CHILD mice presented a similar (2-way RM ANOVA, *p*>0.05,) decrease in calcium-related neuronal activity (-67± 2% from control activity at 20°C; p<0.05, Fig. 6), suggesting that depression of neuronal activity to thermal stimuli remains unaltered after early in life concussion.

**Figure 6.**
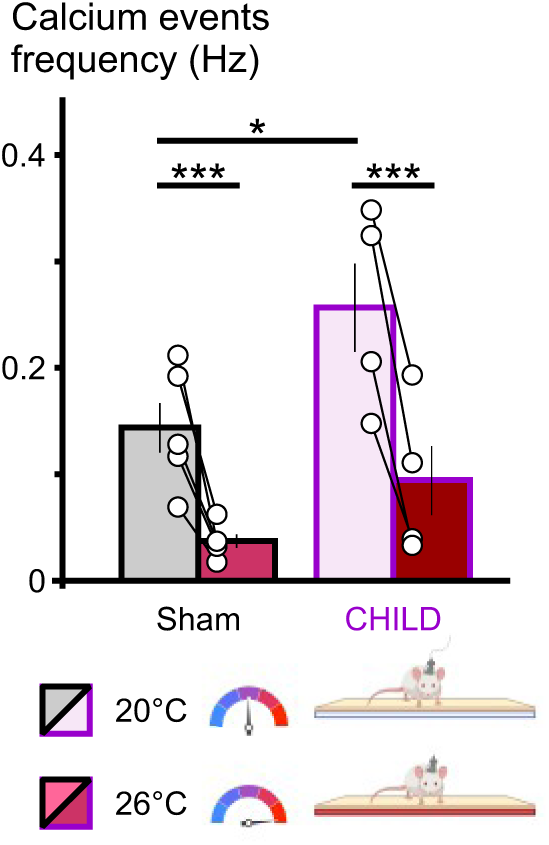
Heat-induced decrease in calcium-related neuronal activity in the SSC remains unaffected by an early life concussion. Group data showing that a 30 seconds exposure to a 6°C increase in floor temperature induced a similar decrease in calcium related neuronal activity in the SSC in both sham (n=5) and CHILD (n=4) mice (2-way RM ANOVA, *F*_(1,17)_=3.339, *p*=0.11). sham mice exhibited a 73 ± 2 % decrease (from 0.14 ± 0.02 Hz to 0.04 ± 0.01 Hz, tukey posthoc, p<0.001, ***) while CHILD mice exhibited a 67 ± 7 % decrease (from 0.26 ± 0.04 Hz to 0.09 ± 0.03 Hz, tukey posthoc, p<0.001, ***). SEM is represented as error bars.

## Discussion

It is well established that traumatic brain injuries are associated with long term alterations in a broad range of physiological functions, even for mild severity, including sensory alterations, contributing to worsening of the patient’s quality of life. To date, the neuronal mechanisms triggered by such events are poorly understood, especially regarding the long-term consequences for which no existing treatment exists for these patients. While children represent the most susceptible population to suffer TBIs, specific research on long-term consequences of those events on the brain that is in development is scarce^42,43^. In our CHILD model, we previously showed long-term behavioral dysfunctions were associated with neuronal loss at 12 months in the hippocampus^18^. However, the neuronal activity has never been assessed over time following the injury in the same individuals. Our novel and timely findings demonstrate for the first time that: (**1**) early in life concussion durably disturbs basal neuronal activity in the SSC at the site of impact, even without overt loss of neurons, (**2**) SSC-neuronal hyperactivity is associated with altered sensory-dependent potentiation of the activity of this network and its related behaviors, (**3**) these alterations are specific to potentiation of neuronal activity since sensory-dependent depression of the activity remains possible, and finally that (**4**) reducing SSC neuronal-activity *via* the potentiation of the endocannabinoid signaling rescues both neuronal plasticity in the SSC and the correlated behaviors. Broadly, we show that the rescue of alterations induced by a concussion during early stages of brain development could be reached a year after injury, which represents a dramatically expanded therapeutic window that is much more amenable for clinical intervention than within minutes after injury, as previously suggested^10–12,44^. Our study also further extends the interest of the endocannabinoid system as a potent and future therapeutic target in the treatment of traumatic brain injuries^10–12,27,34,44–48^ and possibly other brain injuries such as stroke^49^, or neurodegenerative diseases^27,50^.

A crucial feature of traumatic brain injuries is that anyone in its lifetime can suffer one. While severe injuries typically lead to heavy behavioral consequences to the patient, the vast majority of TBI are considered mild. Most mTBI patients do not enter typical hospital emergency rooms and if so are typically quickly discharged with no clinical following. Epidemiologic studies indicate that about 20% of adults suffering mTBIs will eventually develop symptoms lasting 3 months or more^2,3^. When considering the precocity of injury, the incidence of adverse outcomes increases. First, children aged 0 to 4 years old represent one of the two most at-risk age-class^51^. Second, it is estimated that when suffering a mTBI, children more susceptible than adults to developing long lasting consequences^4,5,52,53^. Previous reports have shown in juvenile animal models of severe TBIs, an increased neuronal activity in the ipsilateral somatosensory cortex^16^ and hippocampus^17^ during the sub-acute phase of the injury (*i.e.*, up to a month post injury). Our work demonstrates for the first time that a single, mild, early in life event triggers long-lasting (*i.e.* at least 12 months post-injury) perturbations of brain network activity and related behaviors. Our results strengthen previous observations in mild juvenile brain injury, showing changes in electro-encephalographic activity within gamma frequency at 1 month post-concussion^54^ and memory dysfunction related with a loss of neurons in the hippocampus at 12 months^18^. We also previously observed remote alterations, outside of the central nervous system with long-term cardiac dysfunction^29^. These studies strongly argue for the need for a deeper and more sustained follow-up of younger patients over the long-term. In clinics, traumatic brain injuries during adulthood have been associated to an increased incidence of neurodegenerative disorders including chronic traumatic encephalopathies, Alzheimer’s and Parkinson’s diseases^55^. In Parkinson’s preclinical models, an increased neuronal activity in the amygdala or subtantia nigra have been associated with an increased vulnerability to develop disease-related symptoms^56,57^. Further studies are therefore required to investigate whether the early concussion -induced hyperactivity we report here in the somatosensory cortex could potentially contribute to progression of neurodegenerative disorders. This also raise the importance of long-term studies in regard of early traumatic brain injuries to understand the multiplicity of mechanisms at play when long-term symptoms develop in order to develop *ad hoc* and targeted therapeutic strategies.

We found that neuronal plasticity of SSC neurons was specifically blunted when it requires increasing neuronal activity. Our findings rely on neuronal plasticity of the SSC induced while the animal is behaving in a paradigm that we designed to exhibit a strong sensory component. Taken individually, and without the parallel use of *in vivo* calcium imaging, these tests could lead to different interpretations, but taken together rather indicate altered sensory processing. For instance, we found that mice that entered the open arms of elevated plus maze presented a significant transient increase in neuronal activity in the somatosensory cortex. The amplitude of this form of plasticity was directly inversely correlated to the number of entries in the open arms (Fig. 3e and Fig. 4i). To our knowledge, this is the first report of this behaviorally-induced form of short-term potentiation of neuronal activity. Dogmatically, a change in the number of entries in the open arms in the elevated plus maze is interpreted as a change in the anxiety level of the animal. We propose that after early in life traumatic brain injury the mice in our experimental conditions were not less anxious, but rather were not able to properly integrate a change in environment (i.e. transitioning from closed to open arm), thereby blunting the anxiogenic properties of the edge. Further work is required to determine whether this form of plasticity is related to sensory integration induced by motion and whisking of the animal^58^.

The specific neuronal responses of layers 2/3 but not layer 4 of the S1 somatosensory cortex induced by the novel object has been described previously^41^. We found that after an early in life concussion mice failed to show an increased interest for the novel object and the neuronal plasticity induced by its presence was blunted. Traditional interpretation of these results would suggest an alteration of recognition memory. In the context of our behavioral approach, we rather interpret these results as the consequence of a loss of sensory integration facilitating object recognition.

The SSC has been implicated through sensory integration in the early stages of motor skill acquisition in humans^59–61^. We found that while sham mice were crossing beams in the eponym task, errors in the right hindpaw positioning were associated with transient neuronal responses in the SSC, suggesting specific neuronal error-related sensory integration. In CHILD mice no response was observed while the errors occurred, suggesting an absence of error-related sensory integration and that could impede motor skill acquisition in the task, therefore leading to the increased number of errors made. We also found that neuronal activity in the somatosensory cortex increased in correlation with the speed of the rotarod in the first trial of the test. Interestingly we found that under that experimental condition the level of activity at the beginning of the test (*i.e.* 4 RPM) was predictive of the performance of the mice to stay on the rotarod when speed was increasing. Our novel observations suggest that this neuronal network is able to perform sensory integration in order to adapt locomotion to the rod-speed until reaching a saturated level of activity, at which point the motor command becomes inadequate, leading to the fall of the animal. This would explain the early fall of mice after early in life concussion that present with an already high basal level of neuronal activity. Interestingly, once learned the performance in the task appeared to become independent of somatosensory cortex activity as the correlation was lost on the next test 3 months later (data not shown). As a consequence, sham and CHILD performed equally on this subsequent test. This would argue that during the acquisition phase of the task, performance requires correct sensory integration in the somatosensory cortex, which once learned, the generation of the most appropriate walking behavior relies on other brain structures likely not affected in our model. This interpretation adds to the view that movement-related somatosensory feedback encoded in S1 is important for correct paw positioning and ultimately performance^62^.

Thermal integration on the somatosensory cortex had been previously described^63–65^. The most recent of these studies aimed at deciphering the cellular encoding of focal thermal information, *i.e.* within seconds following the exposure. Our experimental set up did not allow for such time and space resolution. Rather we analyzed neuronal activity over the 30 seconds of exposure to increased temperature of the heating pad and unmasked a depression of the activity of the somatosensory cortex. Such a depression of neuronal activity had been described in rats, with local heat to the scrotal skin^65^. Therefore, the depression of the activity of neurons in the somatosensory cortex we describe here could be attributed to a rebound inhibition following the thermal change or to the overall thermal encoding of all the skin patches exposed on the heating pad (skin patches of the fore and hindpaws, belly, and/or scrotum). In that experimental context, both sham and CHILD mice exhibited similar responses. This would argue for the hypothesis that only sensory integration requiring an increased neuronal activity is altered by an early in life concussion.

Highlighting the need for interpretation of behavioral paradigms in their context rather than according to dogmatic interpretations, we found previously unappreciated alterations of neuronal plasticity in the SSC after early in life concussion, and this lasted up to a year after the concussion. These alterations were specific to increases in neuronal activity as exposure to an increase in 6°C induced a reduction in SSC calcium-related neuronal activity that was similar in both groups. Our hypothesis is that the increased activity either acts as neuronal noise that prevents proper sensory integration (*i.e.* change in inputs) or saturates cellular firing, making impossible further increase in activity with a ceiling effect (*i.e.* change in cellular properties). Our pharmacological manipulation of the endocannabinoid system supports the later hypothesis. Endocannabinoids are critically involved in both synaptic transmission regulation^19,20^ and cell excitability (notably regulating GIRK channels and Ih currents^66–69)^. While both actions would explain the reduction in basal activity following endocannabinoid degradation inhibition, a reduction in synaptic inputs induced by an endocannabinoid tone increase in those conditions could not explain neuronal potentiation rescue, unless though a dis-inhibition mechanism.

Our study also addresses the fundamental question of whether the long-term changes induced by early in life event can be reversed by an intervention that is amenable within a clinical setting. While blocking endocannabinoid degradation within minutes of injury had been proposed as a treatment in more mature preclinical models, relying of the anti-inflammatory properties of endocannabinoids to prevent the development of the primary injury^10–12,44^, we tested such treatment at time from injury when post-concussive symptoms have already developed (*i.e.* a year after concussion). We found that modulating this system according to that protocol could successfully rescue basal neuronal activity and neuronal plasticity in the somatosensory cortex and the associated behaviors. While definitively showing that this rescue was due to the pharmacological agent, the fact that when tested a week later with a vehicle injection, animals exhibited again the post-concussive deficits brings up a limitation of that approach with the question of the treatment efficacy duration. Further work is therefore critical to identify an optimal drug exposure (dose, frequency) that would infer durably a reversal of these post-concussive symptoms. In addition, the endocannabinoid release is tightly linked to neuronal activity^20,70^. Targeting their degradation presents an advantage to firstly regulate the most active neuronal networks, minimally disturbing those with low to normal levels of activity. As a consequence, this strategy is expected to minimize off-target effects.

Sensory integration is a major component of brain processing to allow adaptation to the environment and to ensure survival. One common thought is that events early during brain development have a minimal impact on brain function because of the relatively high plasticity capacity of the immature brain. Clinical studies reveal that pediatric traumatic brain injuries can lead to long-term post-concussive symptoms^4,5^, a persistent increase in basal brain activity associated to blunted plasticity is a previously unappreciated mechanism for the development of long-term alterations of behaviors post-injury (*i.e.* symptoms). Targeting basal neuronal activity could provide an innovative therapeutic strategy to prevent the development of post-concussive symptoms following an early in life concussion. Future work should evaluate the relevance of the blockade of the activity-dependent modulation of endocannabinoid degradation.

## Supporting information

Extended data for Badaut et al 2024

## Acknowledgements.

This research project was funded by Eranet Neuron MISST and Neuvasc (JB), Agence Nationale de la recherche (JB), Nanospace (JB) and NIH, Grant No. (1R01NS119605-01 AO and JB). We thank Richard Rouland and Jonathan Zapata for experimental advice and helpful discussions.

## Author contribution

JB designed the experiments, interpreted the results and proofed the manuscript. LH performed CHILD and all MRI scans. MM performed immunohistochemistry. BPN analyzed MRI scans and tractography. AO designed the experiments, interpreted the results and proofed the manuscript. C.J.D. designed and performed the experiments, the analysis and interpretation of the data and wrote the manuscript.

## Author information

Reprints and permissions information is available at www.nature.com/reprints. The authors declare no competing financial interests. Correspondence and requests for materials should be addressed to CJD.

## Methods

### Animals

CD1 mice were initially obtained from Janvier (Le Genest-Saint-Isle, France) and subsequently bred in–house. Animal were kept in groups, in standard housing conditions (21°C, 55% humidity, 12h light-dark cycle) and had access to *ad libitum* food and water. At postnatal day 17, 9 males were randomly assigned to a sham (n=5) or juvenile mTBI (CHILD, n=4) group. Forty-one to forty-six days later animals from both groups were stereotaxically transfected and implanted with a miniscope lens (Inscopix GRIN lens, Palo Alto, CA). A month later, and every 3 months after the CHILD/sham procedure and up to a year afterwards, animals were tested in behavioral challenges while calcium imaging recordings were performed (Figure 1). All animal procedures were carried out in accordance with the University of Bordeaux animal care committee regulations, French laws governing laboratory animal use (authorization #29324-2021012118549817 v3), the European Council directives (86/609/EEC) and the ARRIVE guidelines.

### Closed-Head Injury with Long-term Disorders (CHILD) model

Concussions were performed at postnatal day 17 as previously described using the CHILD model^18,28^. Briefly, under isoflurane anesthesia (2.5%, 15.l/min) male mice were impacted over the left somatosensory cortex using an electromagnetic impactor (Leica Impact One Stereotaxic impactor, Leica Biosystems, Richmond, IL, USA) with a 3 mm round tip (3 m/s, 3 mm depth and 100 ms dwell time). For the sham procedure, animals underwent the same procedure but were moved away from the impactor before impact. All mice were allowed to recover in an individual cage before being returned to their home cage.

### Surgical procedure

Viral transfection and miniscope implantation were performed at postnatal days 58 to 63 on CHILD and sham mice. Forty-five minutes before the procedure, mice received a subcutaneous injection of buprenorphine (0.05 mg/kg) and saline for hydration. Under isoflurane (Centravet, Mazères, France) anesthesia (induction at 4%, then maintained at 1 to 2 %), mice were then locally shaved and disinfected (betadine, Centravet, Mazères, France). After securing the animal’s head into a stereotaxic frame, a subcutaneous injection of lidocaine was performed and the scalp was removed. Two skull screws were secured on the right part of the skull while a 1x1.4 mm cranial window was drilled over the left somatosensory cortex (AP: -1, ML: -1.8). Viral injections (INSCOPIX ready-to image AAV1.camk2a.GCaMP6f.WPRE.bGHpA) were performed using a Hamilton syringe mounted on a syringe nanoliter infusion system (legato 130, KD scientific, Holliston USA, 250nl/site at 25 nl/min) at the same location (DV: -0.4 and -0.6). After 10 min to allow for viral diffusion, a GRIN lens (Inscopix) was lowered into position (AP: -1, ML: -1.8, DV: -1.2). Brain tissue was then covered with a thin layer of kwik sil (WPI, France) and the grin lens position secured with C&B-Metabond (Parkell, Edgewood, NY, USA) and the skull screws. This was then embedded into an additional layer of dental cement. Mice recovered for at least 3 weeks to allow GCaMP6f expression.

### Behavioral evaluation

All behavioral evaluations were performed together with calcium imaging recording. Therefore, each behavioral session was preceded by the mounting of the miniscope onto the GRIN lens followed by a 5-minute habituation period. All sessions were performed during animal’s day-cycle (around 3 hours after light), in a quiet environment and under controlled lighting. Each apparatus was cleaned with 75% ethanol before testing of each animal to prevent potential bias due to olfactory cues.

### Behavioral evaluation – Elevated plus maze

The homemade apparatus was made of a black polymer and consisted of 4 arms of 35 cm length and 6 cm wide, with 2 opposite arms with 18-cm-high walls (typically 15 lux) while the 2 other arms were opened to the depth (typically 60 lux) and a central square of 6x6 cm. The apparatus was elevated to a height of 60 cm above a platform, itself 60 cm above ground. The animal was positioned at the start of each test on an open arm, facing towards the edge, and was allowed to explore for 5 minutes. The time and distance spent into the open and closed arms, as well as in the central square were monitored and analyzed with Any-Maze (version 7.3, Stoelting Europe, Dublin Ireland). Head entrances in each part of the apparatus were also timed for synchronization with calcium imaging recordings. This test was performed 6, 9 and 12 months after the injury.

### Behavioral evaluation – Novel object recognition test

The apparatus consisted of a custom-made arena (typically 60 lux) with a uniform, smooth black floor (45 x 45 cm) and walls (35 cm high). On day one, mice were habituated to the empty open field during 5 minutes. On days 2 to 4, two standard 50 ml transparent culture flasks (4 cm wide, 11 cm tall, and 2.5 cm wide) with a blue cap filled with red-stained sand identical were positioned in opposite corners, 15 cm away from two consecutive walls. Animals were allowed to explore the arena and the objects for 5 minutes per session. On day 5, one of the objects was switched to a novel object consisting of 2 large white building blocks (8 cm wide, 2 cm tall, and 2.5 cm width) with a smaller light blue building block (6 cm wide, 2cm tall and 2.5 cm width) in between, and mice were allowed to explore for 5 minutes. Tracking was performed using Any-Maze. An area of 5 cm around each object was defined as the area of interest.

The discrimination index was calculated as the percentage of time spent exploring the target divided by total time spent exploring both targets.

### Behavioral evaluation – Beam walk

For this test, mice had to cross a 30 cm custom-made beams of 3, 2 or 1 cm wide (2 trials each) transparent plastic. The starting point was highly illuminated (120 lux) while the goal area was dim (15 lux). Each trial was videotaped for offline recording, focusing on the right side of the animal. Time to cross the beam was manually scored and errors of the rear right paw were precisely timestamped.

### Behavioral evaluation – Rotarod

This test was designed to focus on sensory integration rather than motor learning. For this purpose, initial phase of the test was performed at the fixed speed of 4 rotations per minute (RPM) and consisted of leaving the animal on the rod for a minute total, replacing the animal on the rod in case of a fall. Once the 1 min criteria reached, the animal was allowed to rest for an extra minute. The animal was then replaced on the rod at a starting speed of 4 RPM. The speed was then immediately linearly increased to reach 40 RPM after 2 min. Time and speed at fall were scored, and the session was videotaped.

### Temperature assay – Hot plate

For this test, the animal was allowed to rest on a plate at room temperature (20°C) for 30 seconds. Mice were then positioned on a vivarium-type heating pad (26°C) for another 30 seconds. The session was videotaped for calcium imaging synchronization.

### Pharmacology

12 months post-injury, the effects of endocannabinoid degradation blockade inhibition were tested. For this purpose, a first 5 min home cage session was performed for baseline. The animal was then injected with JZL^184^ (4-nitrophenyl 4-[bis(2H-1,3-benzodioxol-5-yl)(hydroxy)methyl]piperidine-1-carboxylate, 18 mg/kg, Sigma-Aldrich, St. Louis, MO, USA, dissolved in 10 % DMSO, 2% Tween 80 in saline) and subsequently positioned back into the cage. Thirty minutes later, calcium imaging recording was resumed for 5 minutes to evaluate the effect of the drug in absence of significant sensory stimulation. Finally, the animal was tested in the elevated plus maze, as described above.

### Calcium imaging sessions

Recording sessions were performed while the animal’s behavior was also videotaped. Synchrony of the videos was ensured with a LED indicating recording of calcium imaging. Calcium imaging videos (1280 x 800 pixels) were sampled at 20kHz with an exposure time of 50 ms and a gain of 2 using a Nvista3 miniscope system (Inscopix, Palo Alto, CA). Once the microscope was connected to the GRIN lens baseplate, the focal plan of the session was set in order to recover the focal plan determined on the very first imaging session, when possible (the first session aimed at identifying the best focal plane with a maximum number of active cells). Typically, this step lasted around 20 seconds.

### *Ex vivo* analysis

For *post-mortem* analysis, one week after the last behavioral assay, mice were deeply anesthetized (100 mg/kg Ketamine/ 20 mg/kg xylazine) and fixed with an intracardiac perfusion of a 4% PFA solution (in PBS), decapitated and the head was post-fixed for 2 days in a 4% PFA solution (in PBS). After washing the heads with PBS-azide (0.1%), miniscope lenses were carefully removed from the skull and the heads were stored at 4°C in PBS-azide.

### *Ex vivo* analysis – Magnetic resonance imaging (MRI)

Ex vivo MRI was performed in a 7T scanner (Bruker BioSpin, Ettlingen, Germany) on whole brain samples from 10 rodents. The brain was placed into a 50ml Falcon tube, secured to minimize movement and immersed in Fluorinert solution (Synquest Laboratories) to minimize susceptibility artefacts. Imaging parameters were as follows: multi echo T2-weighted imaging (T2WI):TR/TE (6000/6.61 ms), FOV (20 x 20 mm), matrix (128 x 128), slice thickness (0.468 mm), and 25 echoes; diffusion tensor imaging (DTI)—TR/TE (1000/38 ms), slice thickness (0.156 mm), 30 diffusion gradient directions (b = 2000 mT/m), FOV (20 x 15 x 10 mm), with a matrix size of 128 x 96 x 48.

### *Ex vivo* analysis – Immunohistochemistry

Once scanned with MRI, brains were extracted from the skull and coronally sliced using a vibratome (Leica VS 1000, Leica Biosystems, Deer Park, IL). The fifty micron-thick slices were stored in PBS-azide (0.1%). On the day of staining, slices were washed in PBS and incubated in blocking solution (1% BSA, 0.3% Triton X-100 in PBS) for 10 min. Slice were then incubated overnight at 4°C in blocking solution containing a rabbit anti-Parvalbumin (PV) antibody (1:400, cat # PA1933, Thermo Fisher Scientific, Waltham, MA). Slices were then washed and incubated blocking solution containing a secondary antibody donkey anti-rabbit Alexa^568^ diluted to 1:1000 (Abcam, Cambridge, MA) for 90 minutes. After several washes, slices were finally mounted onto glass slides and cover-slipped using Vectashield (Vector Laboratories, Burlingame, CA) with DAPI (1/10000, Thermo Fisher Scientific). Slides were then kept at 4°C until imaging. Fluorescence image acquisitions were performed using a DS-Qi1Mc camera (Nikon Europe, Stroombaan, Netherlands) mounted onto an epifluorescence Nikon eclipse 90i microscope (Nikon Europe) and using the NIS Element software (Nikon, version 4.30.02). LED illumination was provided by a pE-300^white^ CoolLED light source (coolLED, Handover, UK).

### Data analysis – MRI processing

Both T2WI and DTI scans were skull stripped with the segmentation tool from the ITK-SNAP software (version 3.8.0, RRID:SCR_002010)^71^. DTI images underwent denoising using the Adaptive Optimized Non-Local Means (AONLM) filter^72^, followed by eddy current and bias field correction^73^. Tractography was then undertaken by using the somatosensory cortex (SSC) and ventral posterior thalamic nuclei (VPLM) regions from the Australian Mouse Brain Mapping Consortium (AMBMC) atlas^74,75^. These regions were non-linearly registered to each animal’s averaged DTI b0s using Advanced Normalization Tools (ANTs, RRID:SCR_004757). The diffusion data were reconstructed using generalized q-sampling imaging^76^ with a diffusion sampling length ratio of 0.75. The VPLM was seeded and tracts to the SSC were reconstructed with a tracking threshold of 0.016, an angular threshold of 65, and a step size of 0.2. In this study, we tracked projections from the VPLM to the SSC through the caudate putamen, with reference to mouse viral tracing data from the Allen Brain Institute (http://mouse.brain-map.org/, Experiment 100141223). Animals whose tracts did not match the anticipated structure were excluded from tractography analysis, leaving us with 3 sham and 3 TBI subjects. Diffusion tensor metrics fractional anisotropy (FA), axial diffusivity (AxD), mean diffusivity (MD), and radial diffusivity (RD) were extracted from the tract^77^. Additional shape metrics used to characterize additional structural properties, including span, curl, volume, and diameter of the tract were also extracted^78^.

### Data analysis – Behavior

For the elevated plus maze and the novel object recognition test, Any-Maze was used to define the behavior of the animals. The time spent and timing of entries in user-defined areas were collected to define neuronal activity in each compartment. For the beam walk and rotarod tests, the data was manually scored. For all these tests, data was fed into an excel template or python script in order to sort calcium events according to the behavior.

### Data analysis – calcium imaging

First steps of the analysis were performed using Inscopix Data Processing Software (version 1.6.0.3225, Inscopix). For the novel object test and pharmacology experiments videos were concatenated in order to follow neuronal activity of each cell precisely. Briefly, processing included a spatial down sampling (factor 2) and filtering (low cut-off 0.05 pixel^-1^, high cut-off 0.5 pixel^-1^). Motion correction was then applied and the ΔF/F movie (mean frame as reference) was created. A maximum image projection was then created and used to manually draw ROIs. These ROIs were then used to extract fluorescence variations across the ΔF/F movie. These traces were then deconvoluted (model order 1, Spike SNR threshold 3) and calcium-related events detected (event threshold factor 4, event smallest decay time 200 ms). Deconvoluted traces and events parameters were then fed into an excel template or python script for event sorting, traces reconstruction and z-score calculation. For each time point, the z-score was defined as the ΔF/F value of that time point minus the mean of ΔF/F values of the session divided by the standard deviation of ΔF/F values of the session and was calculated with a python script.

### Data analysis – classification algorithm

3 modalities were fed to a home-made classification algorithm using data obtained in neutral condition (home cage, average neuronal activity), amplitude of the network’s response to the entry in the open arms of the elevated plus maze and number of entries of the animal in those arms. Data was converted into numerical arrays for TensorFlow compatibility. A neural network model was constructed using TensorFlow’s Keras API, comprising multiple densely connected layers and an output layer using the softmax activation function for multi-class classification. The model was compiled with the Adam optimizer, categorical cross-entropy loss function, and accuracy as the evaluation metric. The model was trained on the dataset comprising data collected at 6 and 9 MPI on all animals. To avoid overfitting of the data, the model was trained on a maximum of 2000 epochs with an early stopping callback function set at 500 epochs as failsafe. Hence, the algorithm reached a loss of 0.0589 with an accuracy of 1. The model’s performance was then evaluated on separate test datasets collected at 12 MPI, with JZL injection and vehicle injection protocols using the evaluate method. Finally, predictions were generated for both test datasets using the trained model.

### Data analysis – codes and statistics

Behavior and calcium imaging were processed using custom-made excel templates and python scripts, all available upon reasonable requests. Data sets used in this study originated from 2 independent replicates and mice that originated from 3 different litters were randomly assigned to the sham or CHILD group. All values are presented as mean ± SEM. For each statistical analysis normality and equality of the variances were assessed. two-sided tests were used and a *p* value < 0.05 was considered as significant. When needed, 2-way ANOVAs with repeated measurements and *post hoc* Tukey tests were performed. All statistical tests on behavior tests were performed on primary data (not normalized), and statistical tests on calcium imaging were performed on event detection performed on deconvoluted ΔF/F traces. For detailed statistical analysis, see the Supplementary table 1. Data are available upon request from the corresponding author.

