## Extended data for Badaut et al 2024 for "Endocannabinoid-mediated rescue of somatosensory cortex activity, plasticity and related behaviors following an early in life concussion"

**This PDF file includes:**

Extended data Table 1

Extended data Figures 1 – 5

Statistics table

**<u>Extended data table 1.</u> Related to figure 2. MRI tractography metrics.**

| | Sham mice (n=3) | CHILD mice (n=3) | Variation |
|---|---|---|---|
| Number of tracts | 188741 ± 43215 | 156295 ± 43729 | -17 % |
| Mean length (mm) | 3.95 ± 0.15 | 4.49 ± 0.17 | +13 % |
| Span (mm) | 21.07 ± 1.37 | 14.44 ± 0.76 | +16 % |
| VPLM to SSC fiber angle (°) | 65 ± 6 | 54 ± 3 | -17% |

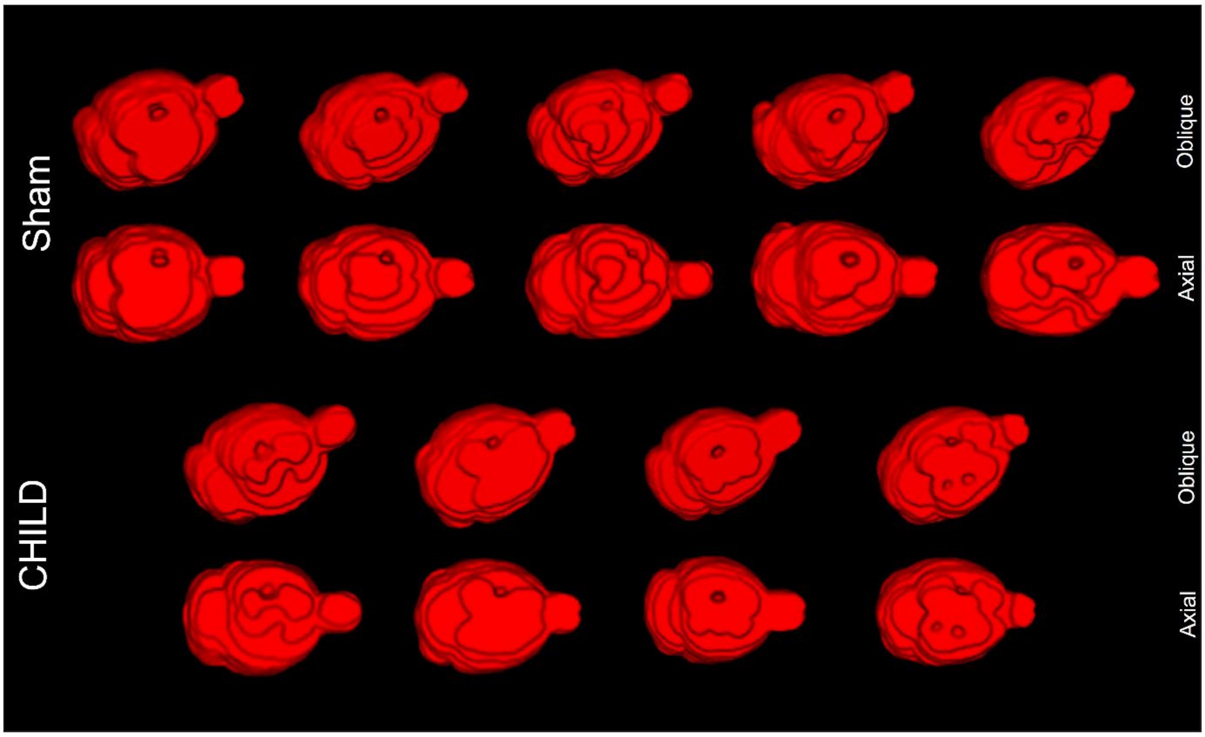


**<u>Extended data figure 1.</u> Related to figure 2. Implantation of the GRIN lenses.**
Twelve months after the traumatic brain injury, brains of sham (n=5, top) and CHILD (n=4, bottom) were fixed and scanned with MRI. 3D-reconstructions allowed for the assessment of the correct location of the lenses (holes).

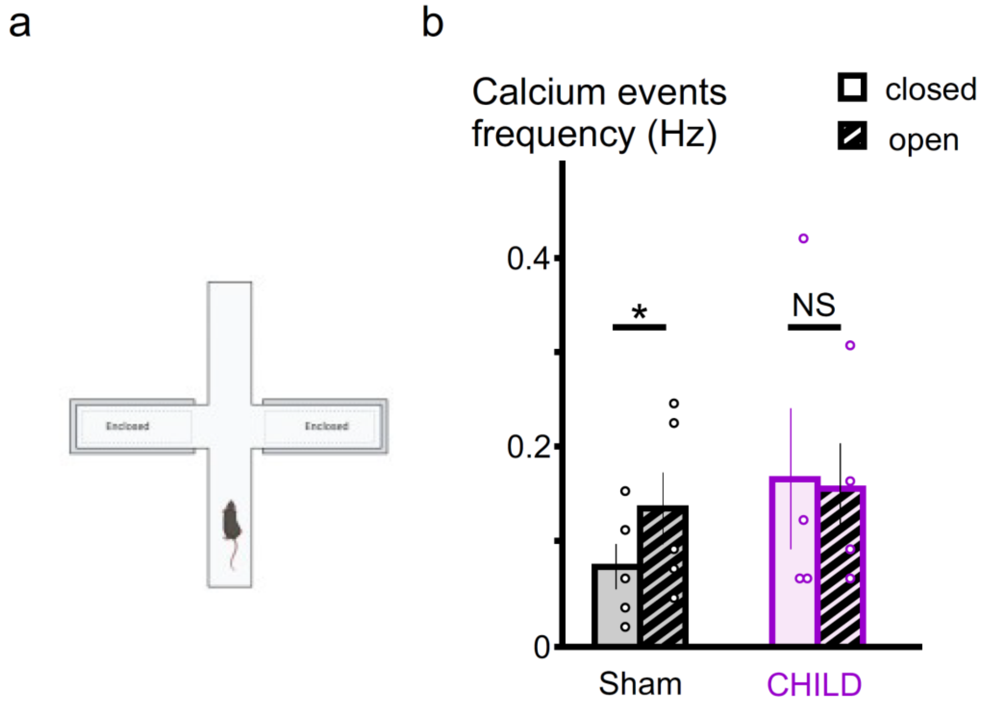


**<u>Extended data figure 2.</u> Related to figure 3. Overall calcium-related neuronal activity in the SSC in the arms of the elevated plus maze.**
**a.** Elevated plus maze **b.** Group data showing that calcium-related neuronal activity in the SSC increased in sham animals (n=5) while in open arms (paired t-test, $p<0.05$), but not in CHILD mice (n=4, paired t-test, $p>0.05$).

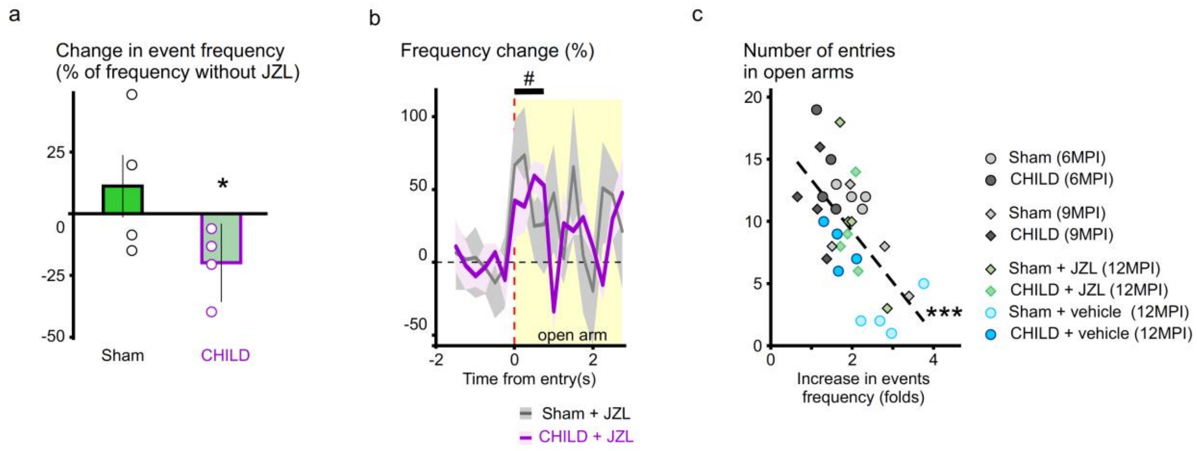


**Extended data figure 3.** **Related to figure 4. JZL184 effects on calcium-related neuronal activity in the SSC in the elevated plus maze.**

**a.** Group data showing the effects of JZL184 (% change in calcium-related events frequency from no drug exposure 30 minutes before) on sham (n=4) and CHILD (n=4) mice in basal condition (typical home-cage). jmTBI mice (n=4) exhibited a significant 26 ± 3% decrease in calcium-related events (paired t-test, $p$=0.027, *), but not sham mice (-7 ± 3%, n=4, paired t-test, $p$=0.2). **b.** Average plots of the averaged normalized calcium-transients frequency in the elevated plus maze in presence of JZL184 of sham mice (grey, n=4) and CHILD mice (purple, n=4). Plots are centered to the entry in the open arms. A similar significant increase in neuronal calcium-related activity occurred in both groups. C. Summary of all scatter plots of the number entries in the open arms plotted *vs* the amplitude of the increase in calcium-transients frequency at the time of entry. There was a significant inverse relationship between these 2 parameters across all experiments (Pearson's correlation, $p$<0.001, ***). Average values are shown as solid lines in **b.**, SEM is represented as the shaded area (error bars in **a.**).

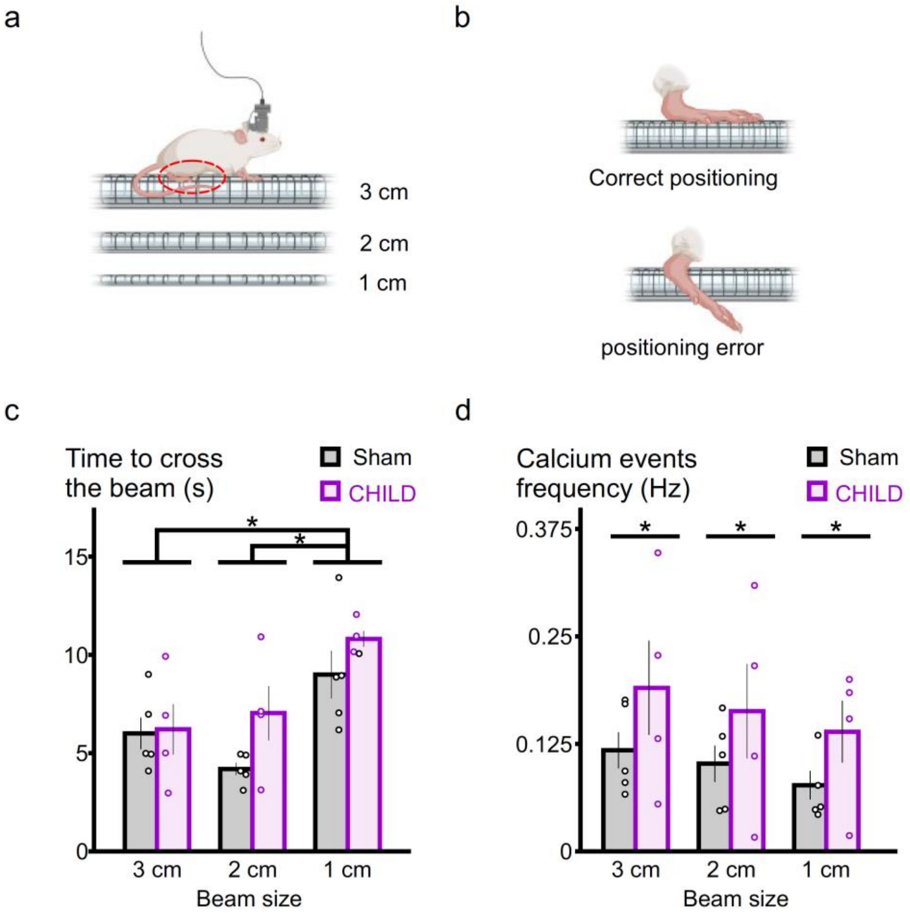


**<u>Extended data figure 4.</u> Related to figure 5. Calcium-related neuronal activity in the SSC in the beam walk task.**

**a.** The beam walk task consisted on crossing beams of a 30 cm length with 3 different cross sections (3, 2 and 1 cm). 2 trials were performed per type of beam. **b.** Schematic highlighting the analysis of the right hindpaw positioning. **c.** Group data showing the time to cross the different beams for sham (n=5) and CHILD animals (n=4). Crossing the beam of 1 cm took more time than for other sizes (2-way RM ANOVA, $F_{(1,26)}$=11.622, $p$=0.001), without difference between groups. **d.** Group data showing the calcium-related neuronal activity in the SSC while sham (n=5) and CHILD (n=4) mice were crossing the beams of different sizes.

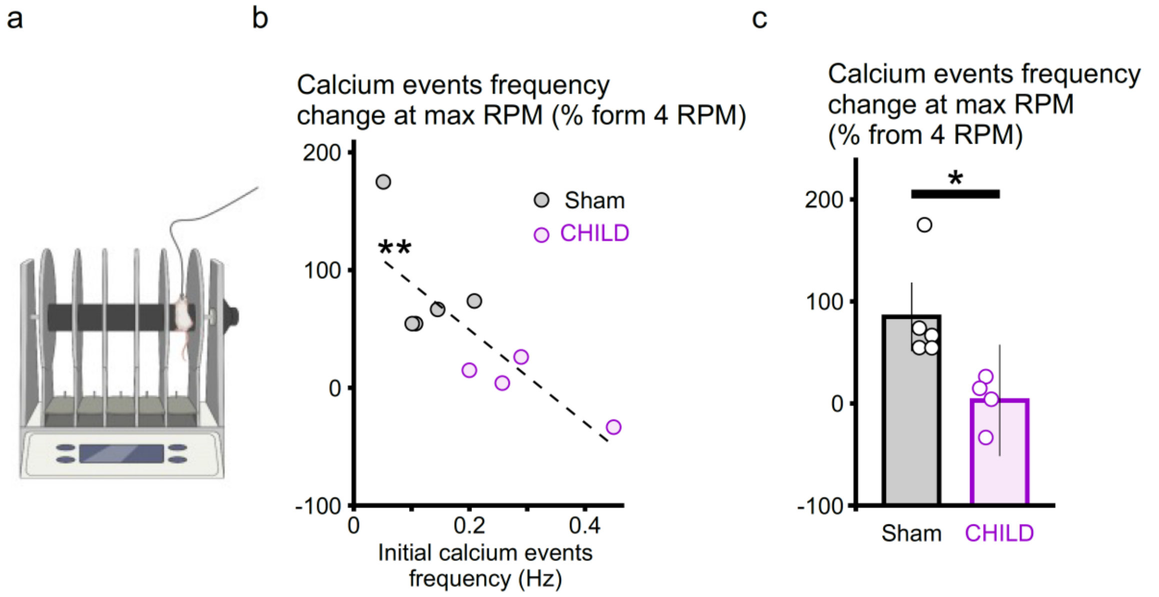


**<u>Extended data figure 5.</u> Related to figure 5. Calcium-related neuronal activity in the SSC in the rotarod paradigm.**

**a.** Rotarod. **b.** plot showing the change in calcium-related neuronal activity from 4 RPM until the fall in sham (n=5) and CHILD (n=4) mice according to their initial calcium-related neuronal activity at 4 RPM. Mice with the lowest initial neuronal activity presented the highest levels of plasticity during the task (Pearson's correlation, $p<0.01$, **). **c.** Group data of the amplitude of the plasticity during the task in sham (n=5) and CHILD mice (n=4). CHILD mice exhibited the lowest plasticity ability (unpaired t-test, $p<0.05$).

**<u>Statistics table.</u>**

Bold P values indicate a significant effect (P<0.05).

| Fig | Parameter analyzed | Conditions | n | Analysis | *F* value | *P* value |
|---|---|---|---|---|---|---|
| in text, relevant to Fig.2c. | calcium events frequency | Sham vs CHILD | 5 - 4 | unpaired t-test | | **0.038** |
| 2c | calcium events frequency | (Sham *vs* CHILD) *vs* (time points) | | 2-WAY RM ANOVA | *F* (4,28)= 0,791 | 0,541 |
| | | Sham *vs* CHILD | 5 - 4 | | *F* (1,28)= 8,064 | **0,025** |
| 3b | time in arms | (open vs closed) in Sham | 5 | paired t-test | | **0,031** |
| | | (open vs closed) in CHILD | 4 | paired t-test | | 0,995 |
| 3d | calcium events frequency | (baseline vs t0) in sham | 5 | paired t-test | | **0,039** |
| | | (baseline vs t0) in CHILD | 4 | paired t-test | | 0,747 |
| 3e | correlation | transcient frequency vs entries open arms | 5 - 4 | Pearson's correlation | | **0,049** |
| 3f | max increase in transcient frequency | (Sham *vs* CHILD) *vs* (time points) | | 2-WAY RM ANOVA | *F* (2,23)= 1,155 | 0,348 |
| | | Sham *vs* CHILD | 5 - 4 | | *F* (1,23)=21,309 | **0,004** |
| 4c | calcium events frequency | (Sham *vs* CHILD) *vs* (ctrl vs JZL) | | 2-WAY RM ANOVA | *F* (1,15)= 10,444 | **0,018** |
| | | (Sham *vs* CHILD) in ctrl | 4 - 4 | tukey *posthoc* | | **0,034** |
| | | (Sham *vs* CHILD) in JZL | 4 - 4 | tukey *posthoc* | | 0,364 |
| | | (ctrl *vs* JZL) in Sham | 4 - 4 | tukey *posthoc* | | 0,450 |
| | | (ctrl *vs* JZL) in CHILD | 4 - 4 | tukey *posthoc* | | **0,002** |
| 4e | number of entries in open arms | Sham vs CHILD | 4 - 4 | unpaired t-test | | 0,814 |
| 4f | calcium events frequency | CHILD (baseline vs t0) in JZL | 4 - 4 | paired t-test | | **0,007** |
| | | CHILD (baseline vs t0) in vehicle | 4 - 4 | paired t-test | | 0,499 |
| 4g | calcium events frequency | vehicle (baseline vs t0) in sham | 4 - 4 | paired t-test | | **0.012** |
| | | vehicle (baseline vs t0) in CHILD | 4 - 4 | paired t-test | | 0.194 |
| in text and supp 3c | correlation | transcient frequency vs entries open arms | 8 | Pearson's correlation | | **<0.001** |
| 4h | correlation | transcient frequency vs entries open arms | 8 | Pearson's correlation | | 0,0883 |
| 4i | correlation | transcient frequency vs entries open arms | 34 | Pearson's correlation | | **<0,001** |
| 5b | Preference index | (Sham *vs* CHILD) *vs* (novelty of object) | 5 - 4 | 2-WAY RM ANOVA | *F* (2,23)= 1,155 | **0,001** |
| | | (novel *vs* habituated) in sham | 5 - 4 | tukey *posthoc* | | **<0,001** |
| | | (novel *vs* habituated) in CHILD | 5 - 4 | tukey *posthoc* | | 0,331 |
| relevant to Fi | Time to cross the beams | (Sham *vs* CHILD) *vs* (beam size) | | 2-WAY RM ANOVA | *F* (2,26)= 0,930 | 0,418 |
| | | Beam size effect | 5 - 4 | | *F* (1,26)= 11,622 | **0,001** |
| | | (Sham vs CHILD) for 3 cm beam | 5 - 4 | tukey *posthoc* | | 0,897 |
| | | (Sham vs CHILD) for 2 cm beam | 5 - 4 | tukey *posthoc* | | 0,084 |
| | | (Sham vs CHILD) for 1 cm beam | 5 - 4 | tukey *posthoc* | | 0,259 |
| 5d | Number of errors | Sham *vs* CHILD | 5 - 4 | unpaired t-test | | **0,042** |
| 5f | calcium events frequency | (Sham *vs* CHILD) *vs* (time points) | 5 - 4 | 2-WAY RM ANOVA | *F* (5,53)= 2,566 | **0,044** |
| | | Sham vs CHILD | 5 - 4 | | *F* (1,53)= 6,913 | **0,034** |
| 5g | Max RPM reached | Sham vs CHILD | 5 - 4 | unpaired t-test | | **0,016** |
| 5h | calcium events frequency | Sham (freq 4RPM *vs* freq max speed) | 5 - 5 | paired t-test | | **0.017** |
| | | CHILD (freq 4RPM *vs* freq max speed) | 4 - 4 | paired t-test | | 0.605 |
| 5i | correlation | initial transcient frequency vs max RPM reached | 5 - 4 | Pearson's correlation | | **0,002** |
| 6 | calcium events frequency | (Sham vs CHILD) vs (temperature) | 5 - 4 | 2-WAY RM ANOVA | *F* (1,17)= 3,339 | 0,11 |
| | | (temperature) in sham | 5 | tukey posthoc | | **<0,001** |
| | | (temperature) in CHILD | 4 | tukey posthoc | | **<0,001** |
